# Engineered Lactiplantibacillus plantarum as a Biosensor Probe for the Lungs

**DOI:** 10.1101/2023.09.27.559779

**Authors:** Michael Brasino, Eli Wagnell, Elise C Manalo, Samuel Drennan, Jared M Fischer, Justin Merritt

## Abstract

Lungs are a frequent site for disease but are difficult to image and probe, potentially exacerbating lung disease, delaying diagnoses and impacting survival. Here, we demonstrate a low cost, minimally invasive method to probe the lungs for disease, using genetically engineered *Lactiplantibacillus plantarum* strain WCFS1. Genetically modified WCFS1 was delivered specifically to the lungs of mice, where it was shown to remain transcriptionally active for several hours and then be cleared without further colonization. WCFS1 was further modified to secrete nanoluciferase as a synthetic biomarker, which traveled from the bacteria in the lung to the urine where it was easily detected. The administered bacteria also secreted nano-luciferase upon detecting a specific peptide secreted by a mouse lung cancer cell line in vitro or in vivo from syngeneic tumors.

## Introduction

There is a growing interest in using genetically engineered bacteria for disease detection. Administration of live bacteria as healthcare has been practiced prior to modern medicine and continues today, such as use of the BCG vaccine.^1^ However, medicinal use of genetically engineered organisms is a recent and exciting possibility. Pioneering work has found that intravenously administered bacteria that naturally infiltrate and multiply within solid tumors may be genetically modified for disease detection and imaging.^2–4^ For example, Shapiro *et al*. engineered tumor-homing bacteria to produce ultrasound contrast by expressing bacterial gas vesicles, these bacteria colonized tumors in vivo, allowing for tumor detection via ultrasound.^5,6^ Others have demonstrated that tumor-targeting bacteria may be delivered non-invasively for some applications. Danino, *et al*. documented the oral delivery of engineered *E. coli* that colonized liver metastases and cleaved intravenously delivered substrates for urinary detection in mice.^7^ Similarly, Riglar et. al. showed that engineered *E. coli* administered orally recorded inflammation as it passed through the mouse gut.^8^ More recently, administration of engineered bacteria to the respiratory tract has been explored with the use of engineered mycobacteria made to suppress pulmonary infections of *Pseudomonas aeruginosa*.^9^ However, the use of engineered bacteria to detect or treat tumors in the respiratory tract has, to our knowledge, not been explored.

Lung cancer kills more Americans than any other cancer. Largely because it is detected at later stages, when treatment options are limited.^10^ The only currently recommended screening strategy is repeated low dose computerized tomography (LDCT) scans, which are costly, insensitive and frequently require individuals to travel long distances to specialized screening centers.^11^ Due to the cost of screening, high false-positive rate, and low sensitivity, LDCT is currently only recommended for individuals with an extensive history of smoking, who still smoke or have quit recently. While these individuals are at significantly increased risk for developing lung cancer, they do not constitute the majority of lung cancer cases, limiting LDCT’s effect on total lung cancer mortality.^12^ Even within this high-risk population, the burdensome nature of LDCT screening has resulted in low uptake, (less than 6% in 2020).^13^ There is a critical need for new lung cancer screening modalities that are more predictive and less burdensome than LDCT scans.

We aimed to explore the possibility of detecting lung cancer through engineered tumor-sensing bacteria, administered directly to the respiratory tract. Unlike previous studies of bacterially mediated cancer detection, we employ a recognized “Generally Regarded as Safe” (GRAS) organism with a well-established history of safe consumption in humans. We have engineered derivative strains as biosensor probes, which do not need to colonize tumors for their detection and can also detect specific tumor secreted molecules *in situ*.

## Results

An initial design consideration was selection of a specific bacterial strain to use as our “chassis organism”, which we would further genetically engineer as a lung cancer probe. To be broadly applicable, engineered bacteria would need to be safe to administer to healthy individuals, with little to no chance of causing infection or adverse effects. Accordingly, we investigated the use of *Lactiplantibacillus plantarum* strain WCFS1 (WCFS1) for this purpose. WCFS1 is a lactic-acid bacteria originally isolated from the saliva of a healthy human child, which has been marketed as a probiotic and has an extensive record of safe consumption in humans.^14,15^ These properties make WCFS1 an ideal candidate for this application. WCFS1 is also genetically tractable, with several plasmid vectors shown to be replicated stably in the species,^16,17^ with well-documented strong promoters and secretion signal peptides,^18–21^ and recently published methods for inserting genetic material into its genome^22,23^. However, while there are many investigations of WCFS1’s interactions with host biology within the gastrointestinal tract, there are relatively few investigations of WCFS1’s effects within the lungs.^24^ Therefore, we aimed to determine the suitability of WCFS1 as a biosensor when specifically delivered to the lower respiratory tract.

### Use of Bioluminescent Strains to Confirm Pulmonary Delivery

We first characterized the gross biodistribution of WCFS1 over time after its administration to the lungs. For this, we used a recombinase-aided homologous recombination method to insert a red-shifted mutant of luciferase from *Renilla reniformis* (*renG*) downstream of the lactate dehydrogenase gene (ldh1) within the WCFS1 genome.^25^ A strong RBS (5’-aggagat) and 9 bp spacer sequence (5’-gtttagaga) was added before the *renG* start codon to facilitate constitutive translation of luciferase. The insertion also contained a downstream erythromycin resistance cassette to facilitate positive selection. Recombinants were selected on MRS plates containing erythromycin, and the desired insertion was verified by amplifying the section of the genome surrounding the insert and Sanger sequencing the PCR product. The resulting strain was designated WCFS1-ldh1-renG, and was found to luminesce with the luciferase substrate coelenterazine (CTZ) when grown in standard laboratory growth media. We then used this luminescent strain to determine the optimal method of delivering suspensions of live *L. plantarum* specifically to the lungs of mice. WCFS1-ldh1-renG was grown to mid-log, centrifuged and resuspended in PBS before being administered. Oropharyngeal administration (OPA) was used to deliver the bacteria specifically to the lungs. Healthy female mice were administered approximately 8 × 10^8^ CFU of WCFS1-ldh1-renG via OPA, then injected with the luciferase substrate coelenterazine-h via tail vein and imaged immediately in a biophotonic imaging system. As shown in **Figure 1A**, luminescence is visible exclusively in the lungs when mice are imaged immediately after administration, indicating that OPA is an effective method for delivery. After 24 hours, coelenterazine-h injection and biophotonic imaging was repeated and no significant luminescence was observed within the lungs, or any other organ, indicating successful clearance and a lack of colonization of WCFS1. By comparison, nebulization failed to produce significant luminescence signal in the lungs and intranasal administration resulted in strong signal only in the area of the nasal cavity **(Supplementary Figure S1A)**.

**Figure 1:**
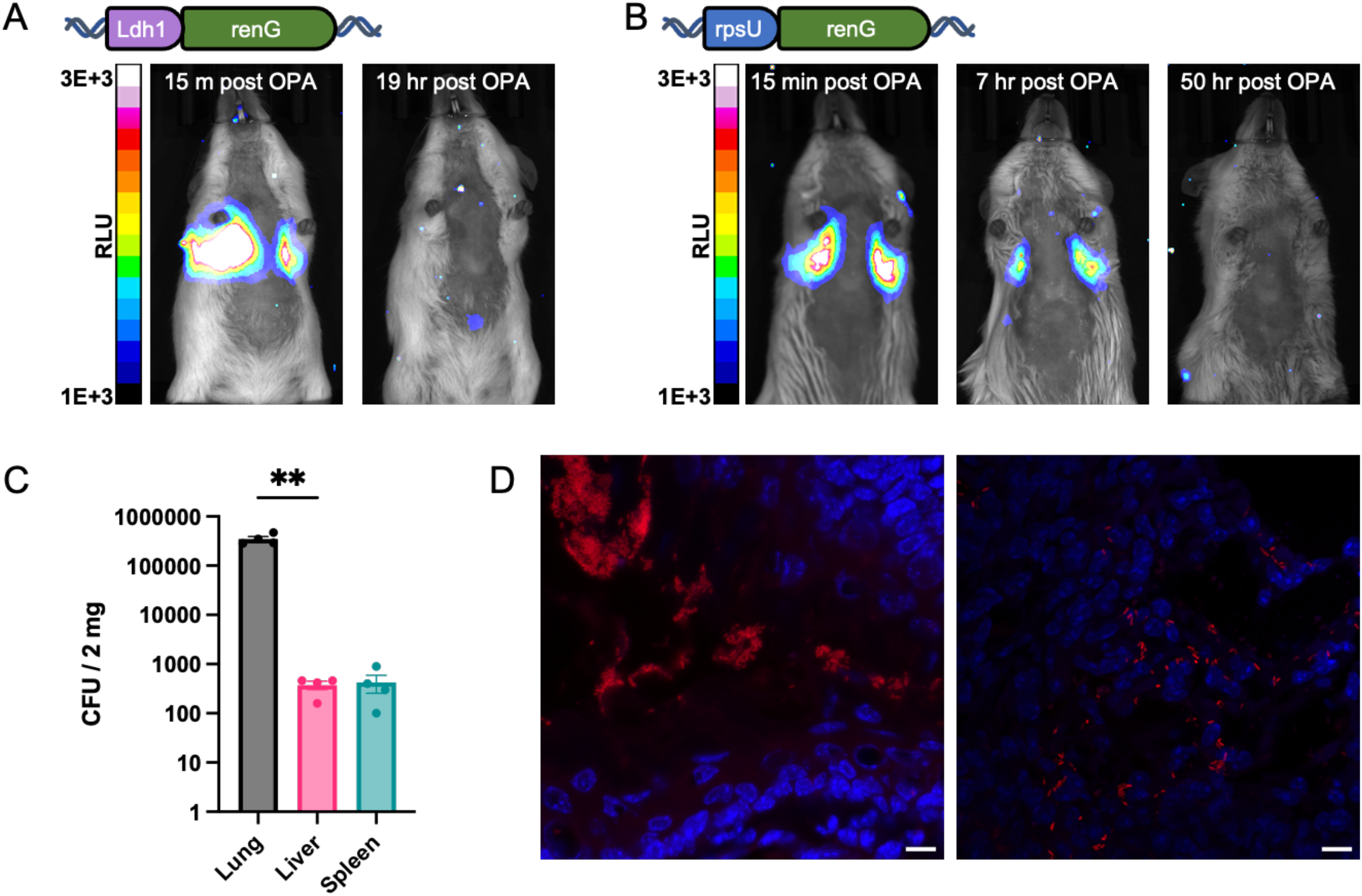
Biodistribution of bioluminescent WCFS1 after pulmonary administration. (**A)** Representative (N=3) in vivo bioluminescence images of the same mouse at the indicated time points after being administered 2 × 10^9^ CFU of WCFS1-ldh1-renG via OPA. **(B)** Representative (N=3) in vivo bioluminescence images of the same mouse at the indicated time points after being administered 7.5 × 10^9^ CFU WCFS1-rpsU-renG via OPA. **(C)** Four mice were administered 4.4 × 10^**8**^ CFU of WCFS1 and then sacrificed and dissected after 6 hours. The indicated organs were homogenized and plated on selective media. CFU’s from 2mg of each tissue is shown. No CFUs were observed from mice not administered WCFS1. **(D)** Micrographs of a mouse lung immediately after administering WCFS1 via OPA. Lungs were cryo-sectioned and hybridized with an oligo probe specific for WCFS1 16S RNA conjugated with Cy5, then mounted in DAPI containing medium. Composite images with DAPI signal in blue and Cy5 signal in red are shown. Micrographs are of main air ways (left) and lower lung (right). Scale bars are both 10 μm.

To confirm WCFS1 distribution, independently of ldh1 expression, we created an additional bioluminescent strain by inserting the same *renG* luciferase downstream of the ribosomal subunit protein S21 (*rpsU*) gene. We also reasoned that *rpsU* expression may be better insulated from fluctuations in bacterial metabolism. This new strain (WCFS1-rpsU-renG) produced slightly lower luminescence when grown in standard laboratory media (**Supplementary Figure S1B)** but still produced a robust and consistent signal.

WCFS1-rpsU-renG was grown to mid-logarithmic phase, centrifuged, and resuspended in PBS, before being administered to healthy female mice via OPA as before. Mice were then imaged immediately and then periodically after in a biophotonic imaging system as before. As shown in **Figure 1B**, bacteria were again delivered specifically to the lungs. After 7 hours, bioluminescence is decreased but remains exclusively in the lungs. And again, no bioluminescence is seen at later time points, indicating a lack of WCFS1 colonization.

Crucially, this pulmonary signal at 7 hours indicated that WCFS1 may be retained in the lungs long enough for sensing activities. Prolonged retention was corroborated by the results of a later experiment shown in **Figure 1C**, in which mice were administered a similar quantity of erythromycin-resistant WCFS1 via OPA, then sacrificed after 6 hours. Mouse organs were then homogenized, diluted, and plated on agar media to determine the number of viable WCFS1 present in each organ. There was over 1,000-fold more CFU/mg in the homogenized lungs compared to the liver or spleen (RM ANOVA F (1, 3) = 63.6, lung vs liver P = 0.0084, lung vs spleen P = 0.0085, liver vs spleen P = .9568) suggesting that the vast majority of the administered bacteria remain in the lung up to 6 hours. Immediately after OPA, mice recover from anesthesia and show no visible signs of distress. After 2 to 4 hours mice became lethargic, but recover by 16 hours. Lethargy was not observed in control mice given an OPA of PBS alone. This response may be caused by an innate immune response to the high bacterial dosage needed for effective *in vivo* imaging in the lungs.

Finally, to gauge the microscopic distribution of WCFS1 in the lungs immediately after OPA delivery, mice were sacrificed and their lungs removed, sectioned and probed for bacteria using fluorescence in-situ hybridization (FISH). For this, a FISH probe (5’-CCAATCAATACCAGAGTTCG) specific for the 16S RNA of WCFS1 was designed and used. The probe sequence was checked for specificity using BLASTN and did not produce significant signal against other non-target bacteria, including similar probiotics within the previous *Lactobacillus* genus (**Supplementary Figure S2**). Slides were additionally counterstained with DAPI to visualize lung tissue. Representative micrographs in **Figure 1D** show WCFS1 (red) clustered in large groups in the main airways, but also well dispersed throughout the lung tissue, and present in all lung lobes.

### Development of Nanoluciferase as a Synthetic Biomarker

We next sought to develop a method for engineered bacteria to signal the presence of disease from within the lungs. To be used in screening applications, an ideal signaling system should be easily measured without complex imaging instrumentation or laboratory services. Consequently, we focused on developing ways for bacteria retained within the lungs to create signals which could be analyzed in host urine. Inspired by synthetic biomarkers used in engineered cell prognostics, we employed a secreted nanoluciferase (nLuc) as a small, easily-expressed, and sensitive synthetic biomarker.^26^ We first confirmed our ability to measure purified nLuc in various media, including murine urine. Urine pH is known to vary widely between individuals and within individuals over time. Thus, to prevent pH variation or other urinary components from interfering with nLuc activity, urine samples were stored at 4°C for no more than 24 hours and then diluted into a 1M Tris-HCl buffered to pH 8.0 before reading. With this approach, purified nLuc was accurately quantifiable in mouse urine down to 2 fg/μl and showed no drop in signal when stored (**Supplementary Figure S3B and C**). We then confirmed WCFS1’s ability to secrete active nLuc. Unlike in mammalian cells, nLuc was internalized and not secreted when expressed in WCFS1 alone (**Supplementary Figure S4A)**. However, secretion could be accomplished by appending a signal peptide from the WCFS1 trans-glycosylase Lp_3050 (RefSeq WP_011102021), which was previously shown to secrete heterologous proteins.^27^ This secreted form of nLuc was then inserted downstream of the *rpsU* gene within the WCFS1 genome, similarly to *renG* above. The resulting recombinant strain (WCFS1-nLuc) was sequence verified and constitutively secreted full length nLuc, as evidenced by western blot (**Supplementary Figure S4B**). This strain secreted approximately 46.1±1.6 ag/CFU of nLuc in 3 hours in DMEM-FBS while secretion was lower, at 19.6±1.6 ag/CFU in PBS with 1% w/v BSA, and 15.5±0.9 ag/CFU in a commercial artificial saliva containing mucin (**Supplementary Figure S4C**).

We then administered mid-logarithmic phase WCFS1-nLuc to female mice via OPA as with the bioluminescent strains above and measured urinary nLuc activity immediately before administration and approximately every 2 hours thereafter. As shown in **Figure 2A**, nLuc signal was detected in mouse urine 2 hours after administration via OPA and increased until 6 hours later at the last time point of the day. The nLuc concentration is lower the next day, and returned to nearly background levels after 50 hours.

**Figure 3:**
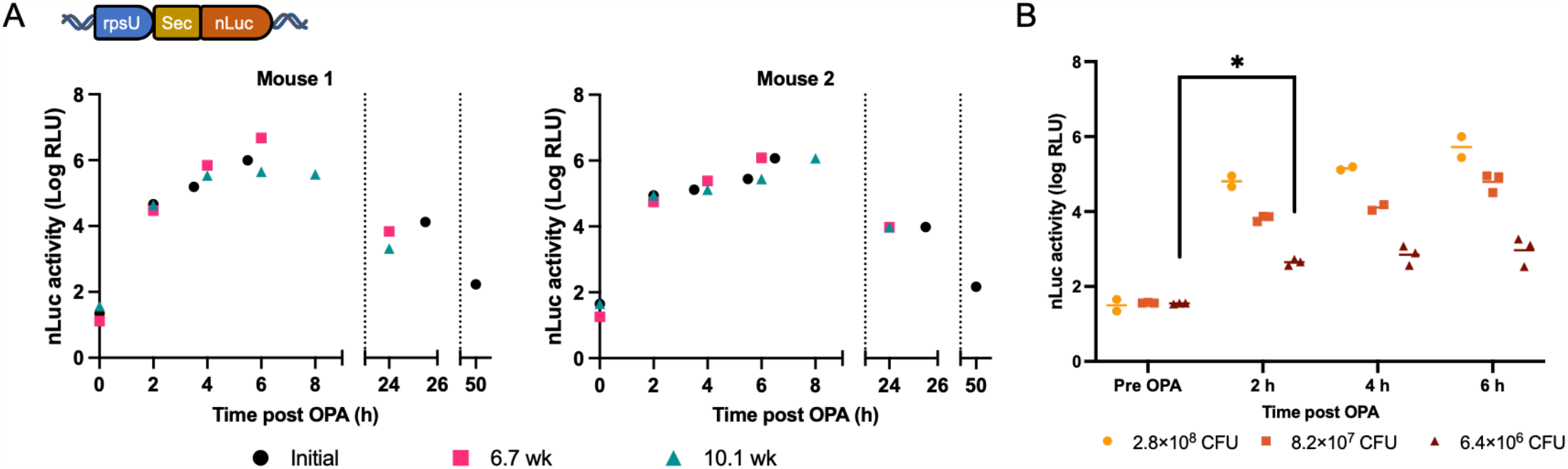
Secretion of nLuc into Urine from WCFS1 in Lungs. **(A)** Urinary nLuc activity from two mice administered 2.8 × 10^8^ CFU of WCFS1-rpsU-nLuc initially, then 8.4 × 10^8^ CFU at 6.7 weeks and 8.0 × 10^8^ at 10.1 weeks. **(B)** Urinary nLuc activity from mice administered the indicated amounts of WCFS1-rpsU-nLuc. Data for 2.8 × 10^8^ CFU are the initial administrations to Mouse 1 and 2 shown in (A). Data for lower dosages are from three separate mice each. Horizontal bars indicate mean signal for each group.

Importantly, the period of high urinary signal immediately after administration corresponds to when bacteria are localized in the lungs, as indicated by the above bioluminescence imaging studies. This suggests that active nLuc enzyme is capable of travelling from the lungs to the urine. However, further studies will be needed to determine if nLuc crosses the pulmonary mucus layer alone, or is transported through some other process, such as phagocytosis of bacteria by alveolar macrophages. Next, we were curious to explore the possibility of using nLuc secreting WCFS1 for longitudinal monitoring of pulmonary disease. To this end, we repeated the OPA administration of WCFS1-nLuc on two additional dates, leaving at least 3 weeks in between each administration to allow for WCFS1 to potentially colonize their hosts or a secondary immune response to potentially develop to either the WCFS1 or nLuc. As shown in **Figure 2A**, there was no obvious drop in urinary signals after these later administrations compared to the initial administration. This indicated that if a secondary immune response did develop, it did not hinder pulmonary WCFS1 from creating urinary nLuc signals. Urinary nLuc signals also returned to their initial baseline before each new administration, suggesting that there was minimal or no colonization with WCFS1.

As urinary nLuc concentrations were well above our limit of detection, we then administered lower dosages of WCFS1-nLuc and again measured urine for activity. Urinary nLuc luminescence is shown in **Figure 2B**. Even with the lowest dosage tested (6.4 × 10^6^ CFU), average urinary nLuc activity at 2 hours post administration corresponded to 56 fg/μl, over 10x above background signals and significantly different from signals pre-administration (Paired t-test, t = 22.93, df = 2, P = 0.0019). Encouragingly, mice administered this lower dosage did not exhibit the acute lethargy observed with higher bacterial dosages, indicating that lower bacterial load may avoid this response.

### Detecting Xylose in the Lungs

To be used as an *in vivo* biosensor, it was crucial to confirm nLuc could be secreted by bacteria within the lungs, in response to conditions there. WCFS1-nLuc growth media was removed before administration, limiting the amount of residual extracellular nLuc administered. However, there remained the possibility that urinary signal originated from residual nLuc expression within the bacteria and that bacteria did not remain transcriptionally active within the lungs. To examine this possibility, we placed the secreted nLuc gene under the control of a promoter regulated by a xylose inhibited repressor protein *xylR*, which was constitutively expressed in the opposite direction upstream. This entire xylose inducible expression cassette was inserted downstream of the *rpsU* gene such that the stop codon for *rpsU* abutted the stop codon of *xylR*, separated by a strong transcriptional terminator. Read through transcription from *rpsU* to secreted nLuc was alleviated by a transcriptional termination sequence placed immediately upstream of the xylose inducible promoter.

As shown in **Figure 3A**, the resulting strain (WCFS1-xylR) showed increased nLuc secretion in a xylose dose-dependent manner, with a 26-fold signal increase and EC50 of 0.23 mg/ml of xylose. We prepared this strain for administration by growing to mid-logarithmic phase and then concentrating in PBS. Then, we administered 6 × 10^8^ CFU of WCFS1-xylR to mice via OPA, either with or without xylose added immediately beforehand. In this way, bacteria were induced with xylose while in the mouse lungs and increased urinary nLuc activity between mice with induced vs. uninduced bacteria could be attributed to bacteria sensing xylose in the lungs and increasing expression of nLuc *in vivo*. As shown in **Figure 3B**, there was indeed a significant increase in mean urinary nLuc activity at 1 h, 3 h, and 6 h post OPA in the four mice administered bacteria with xylose vs. the four mice administered bacteria alone (Multiple paired t tests, [t-ratio, df, P] = [5.467, 6, 0.0015], [3.819, 6, 0.0087] and [3.514, 5, 0.0170] respectively), indicating that bacteria remain sensitive to their environment, and transcriptionally active post administration to the lungs. At 6 hours post OPA average urinary nLuc concentrations were over 500-fold higher in xylose administered mice than in controls (26.4 pg/μl vs 49.7 fg/μl). Average signal remained higher for mice administered xylose until the next day, though barely lost statistical significance, likely due to heterogeneity between individual mice ([t-ratio, df, P] = [2.345, 6, P 0.0574]).

**Figure 3.**
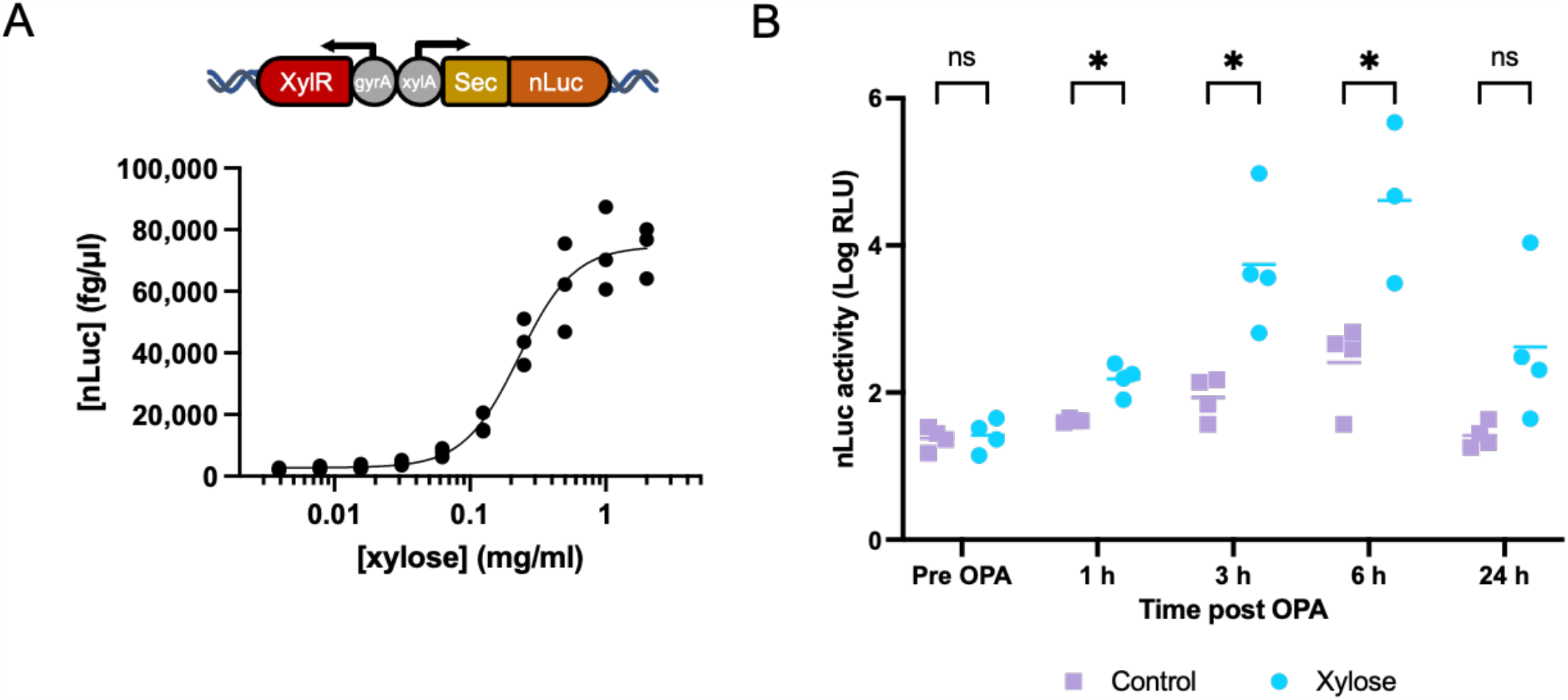
**(A)**nLuc concentrations produced 3 hours after inducing mid-log WCFS1-xylR with DMEM-FBS containing the indicated concentration of Xylose. Data represent the average of 3 biological replicates. **(B)** WCFS1-xylR was grown to mid-log, concentrated in PBS and mixed 1:1 with a solution of 20% w/v xylose/saline or saline alone immediately before OPA to four mice each. Urinary nLuc activity of each mouse was measured at the indicated times after administration. Individual signals from all eight mice at each time point are shown, other than at 6 h, when we failed to collect urine from a single mouse administered xylose. Averages for each group are indicated as horizontal bars. Individual un-paired T-tests (two tailed) were used to compare signals from xylose and control mice at each time point. P values are (from left to right) 0.769, 0.0015, 0.0087, 0.0170 and 0.0574 for pre OPA, 1 h, 3 h, 6 h, and 24 h, respectively. Significance (*) is indicated for P < 0.05.

### Model Tumor Detection with Peptide-Sensing Bacteria

We next sought to model how bacteria might sense a specific tumor biomarker within the lungs and respond by secreting nLuc as a synthetic biomarker. For this we leveraged the high-specificity and sensitivity of intercellular peptide signaling receptors common in Gram-positive bacteria like WCFS1. These receptors are capable of detecting sub-nanomolar concentrations of specific short peptides, allowing for excellent sensitivity and specificity. To model how one could leverage this exquisite specificity to identify tumors, we modified a mouse lung cancer cell line (Lewis Lung Carcinoma) LLC to secrete the short bacterial peptide SppIP, and engineered WCFS1 to secrete nLuc in response to SppIP via the two-component receptor system SppKR. As shown in **Figure 4A**, the SppIP peptide acts as a model tumor biomarker which is specifically secreted only by tumor cells and not normal tissue.

**Figure 4:**
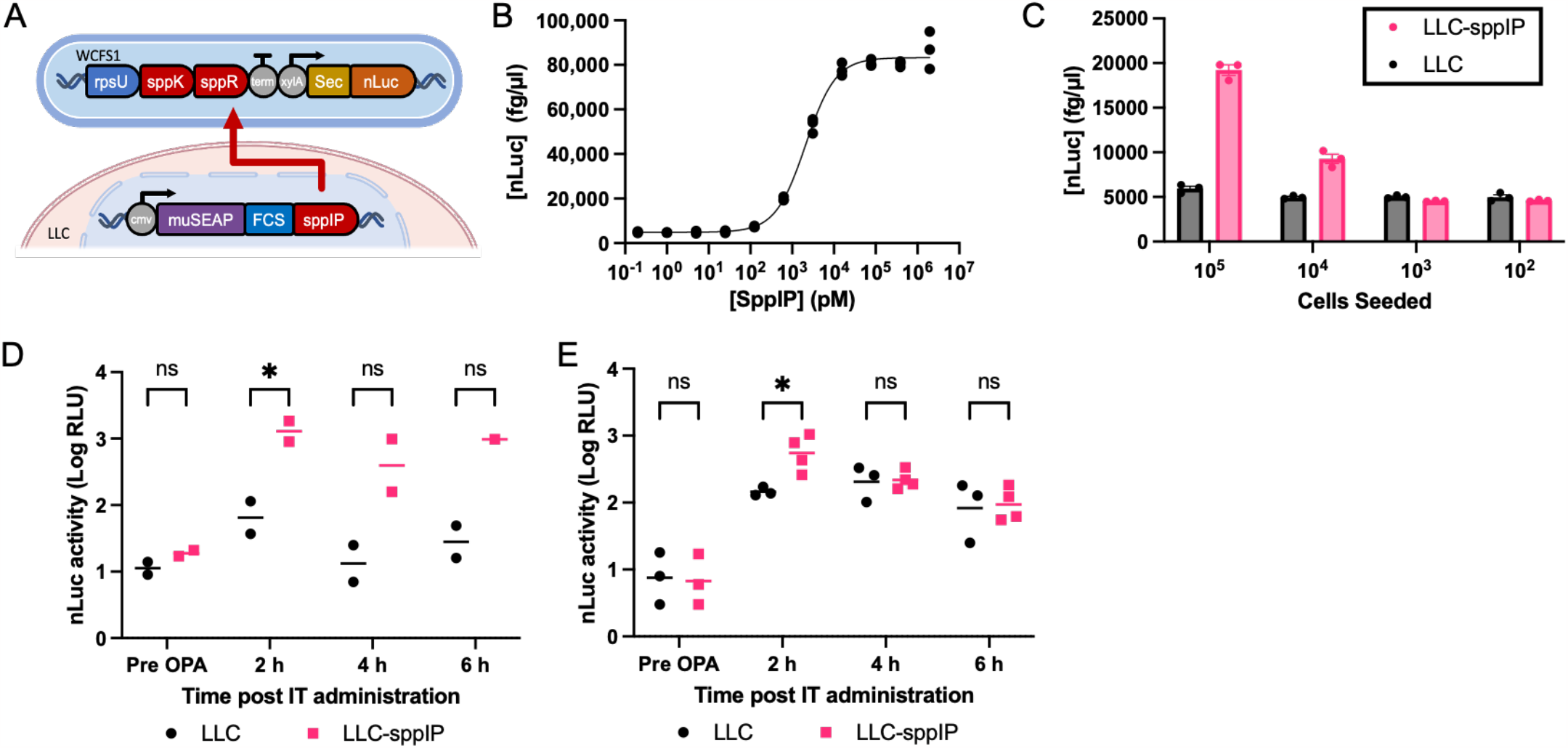
Bacterial Detection of Tumor-Secreted Peptide. **(A)** Schematic of the gene circuits used to secrete SppIP peptide in Lewis Lung Carcinoma cells (LLC) and to detect SppIP in WCFS1. SppIP is secreted by genetic fusion with murine secreted embryonic alkaline phosphatase (muSEAP) via a furin cleavable linker sequence (FCS). In the WCFS1-sppKR strain, sppIP activates a two component sensor system, which in turn drives secreted nLuc expression. **(B)** The concentration of nLuc secreted from WCFS1-sppKR bacteria after incubating with various concentrations of synthetic SppIP peptide in DMEM-FBS for 3 hours at 37ºC. Data from 3 biological replicates are shown along with the best fit dose response curve (Hill equation with variable slope). **(C)** The concentration of nLuc produced by WCFS1-sppKR from cell media 24 hours after seeding the indicated number of peptide producing (LLC-sppIP) or un-transfected (LLC) cells in 200 μl in a 96 well plate. Bar height represent average of 3 cell culture replicates and error bars are S.E.M. **(D)** Mice were injected subcutaneously with either LLC or LLC-sppIP. After two weeks subcutaneous tumors had developed. Two mice from each group were then injected intratumorally with WCFS1-sppKR resuspended in PBS. **(E)** After 5 additional days, all mice (3 with LLC tumors and 4 with LLC-sppIP tumors) including the two previously injected mice from each group, were injected with WCFS1-sppKR resuspended in PBS. In (D) and (E) Urinary nLuc activity for each mouse is shown and horizontal bars represent group averages. Individual unpaired T-tests (two tailed) were used to compare signals from LLC and LLC-sppIP injected mice at each time point. P values from left to right across (D) and (E) are: 0.1673, 0.0462, 0.0925, 0.1694, 0.8814, 0.0160, 0.8728 and 0.8515. Significance (*) is indicated for P < 0.05.

LLC cells were transfected with a lentiviral vector containing the murine secreted embryonic alkaline phosphatase (muSEAP) cDNA sequence fused to a mouse codon-optimized coding sequence for the SppIP peptide at its c-terminus. Between these fused sequences, we inserted a furin cleavage site (TRSRKKR) and expressed the entire construct from a strong constitutive CMV promoter. The highly secreted muSEAP acted as a carrier protein which transported the SppIP peptide extracellularly. There, furin protease which is ubiquitous in murine tissues and secreted by LLC cells *in vitro*, cleaved SppIP from muSEAP, freeing it to be detected by engineered bacteria. This transfected LLC cell line was designated LLC-sppIP. WCFS1 was engineered to detect SppIP by inserting the SppKR two component receptor and nLuc behind the constitutive *rpsU* gene as before. This was done such that *sppK* and *sppR* were co-transcribed with *rpsU*, but transcriptionally insulated from the downstream transcript for nLuc by a pair of strong transcription termination sequences. The coding sequence for nLuc was placed behind the promoter for *sppA*, which is activated by SppK and R in the presence of sppIP. The resulting strain was designated WCFS1-sppKR. As shown in **Figure 4B**, WCFS1-sppKR could detect as little as 125 pM of synthetic SppIP peptide diluted in complete cell culture media (t-test, [t, dF, P] = [9.240, 4, 0.0008]). When maximally induced, WCFS1-sppKR produced a nLuc signal 17-fold higher than without SppIP. Importantly, media taken from LLC-sppIP cells grown *in vitro* could activate secreted nLuc expression in WCFS1-sppKR, indicating that LLC-sppIP cells successfully secreted and processed the sppIP peptide (**Figure 4C**). As expected, and the amount of activation trended with the number of LLC-sppIP cells originally seeded, as more secreting cells led to a higher concentration of sppIP in the surrounding media. WCFS1-sppKR produced significantly more nLuc in media from LLC-sppIP cells compared to LLC cells, even when as little as 10^4^ cells were seeded (P = 0.0013). WCFS1-sppKR was also specific to the sppIP peptide, as media from LLC cells secreting muSEAP fused to alternative peptides failed to activate increased nLuc secretion (**Supplementary Figure S5**). This indicated that WCFS1-sppKR bacteria was capable of detecting tumor secreted peptides *in vitro*.

We next determined whether WCFS1-sppKR was capable of detecting SppIP secreting tumors *in vivo*. For this, we subcutaneously injected **5 × 10**^**5**^ LLC-sppIP or LLC cells into 4 and 3 mice (respectively), and allowed tumors to develop from these cells. After 2 weeks, tumors had visibly grown and urine from mice bearing LLC-sppIP tumors contained SEAP activity, while mice injected with LLC cells did not (**Supplementary Figure S6**). We then injected 4 × 10^8^ CFU of WCFS1-sppKR bacteria intratumorally into two mice from each group and collected urine every 2 hours after. As shown in **Figure 4D**, the urine collected at the first time point (2 hours post-injection) gave the highest nLuc signal of all samples. As expected, the urinary nLuc signal from mice bearing sppIP secreting tumors was significantly higher than those with LLC cells alone (t test, [t-ratio, df, P] = [4.492, 2, P = 0.0462]). After an additional 5 days of tumor growth, all mice (including the two from each group previously administered bacteria) were again injected with WCFS1-sppKR and urine measured for nLuc activity as before (**Figure 4E**). Again, nLuc activity was significantly higher in LLC-sppIP vs. LLC bearing mice 2 h post administration (t test, [t-ratio, df, P] = [3.570, 5, 0.0162]). These results indicated that the WCFS1-sppKR could detect tumor secreted peptides *in vivo* and respond by secreting nLuc into mouse urine. Interestingly, the nLuc signals were significantly lower than when similar dosages of nLuc secreting bacteria were administered via OPA, with the mean signals observed at the first 2 h collection corresponding to only 200 fg/μl and decreasing thereafter. Also unlike OPA, no acute toxicity or immune response was observed in any mouse after bacterial injections.

## Discussion

Here, we have explored the use of a genetically engineered probiotic bacteria for tumor detection in the lower respiratory tract. In the envisioned screening strategy, engineered commensal bacteria would be aspirated by individuals at risk of developing lung cancer, which could be performed with either a nebulizer or possibly by simply eating a carrier food such as yogurt. Once distributed over the mucus membrane above an early lung tumor, engineered bacteria would detect specific peptides present within the tumor microenvironment and secrete synthetic biomarkers in response. In this way, engineered bacteria would amplify a weak localized signal within the tumor microenvironment into a robust systemic signal easily detected in accessible fluids such as urine.

Our results delivering the probiotic *Lactiplantibacillus plantarum* WCFS1 via oropharyngeal administration indicate the vast majority of these bacteria remain in the lungs for several hours after delivery. While high dosages of these bacteria likely produced an innate immune response in mice, we did not observe any evidence of a secondary immune response, allowing for repeated administrations without worsening symptoms. Furthermore, lower dosages of these bacteria were still capable of producing robust urinary signals from within the lungs. This negative response to high dosages may be partly explained by the incompatibility of the human commensal strain WCFS1 with mice. While the WCFS1 strain may be feasibly translated to humans, it’s tolerance by mice is less established. Others have genetically engineered *Escherichia coli* strains, which are commensal in mice to study their utility in the murine gut^8,28^ Similarly, the use of alternative strains of *Lactiplantibacillus* or other lactic acid bacteria which are commensal in mice might alleviate this innate response, and facilitate studies in the murine respiratory system. In the envisioned application to lung cancer detection, bacterial dosage will ultimately be constrained by the sensitivity of these bacteria for cancer biomarkers, and the prevalence of these biomarkers at early stages of disease.

We also demonstrate the feasibility of detecting synthetic biomarkers in urine secreted by WCFS1, resident in the lungs. Using nanoluciferase (nLuc) as a synthetic biomarker, we observed strong urinary signals as early as two hours after pulmonary administration which persisted for at least 6 hours. Crucially, this is when our biodistribution studies show that the WCFS1 remain largely in the lungs. Repeated administrations of nLuc secreting WCFS1 also provided consistent signals, indicating that any secondary immune response to the bacteria or nLuc does not interfere with nLuc’s ability to reach the urine. Furthermore, our results using a strain which secretes nLuc in response to xylose indicate that WCFS1 remains transcriptionally active and capable of induction in the lungs. Interestingly, the ratio of nLuc concentrations from induced vs. uninduced conditions was lower *in vitro* than when bacteria were induced in the lungs and nLuc measured in urine. This difference might be attributed to some consistent degradation or clearance of nLuc *in vivo* which is overwhelmed under inducing conditions. Protein engineering approaches such as the attachment of hydrophilic tags might alleviate this degradation and further increase the accessibility of nLuc in urine.^29^

Finally, we have shown that WCFS1 is capable of detecting tumor secreted peptides *in vivo*, which underscores its suitability for detecting specific tumor biomarkers. Previous efforts in the field have developed transcriptional sensors for hallmarks of the tumor microenvironment, such as low pH, hypoxia, and elevated lactate concentrations.^30^ However, these sensors have been built to function in *E. coli* and primarily Gram-negative species and not immediately suited to lactic acid bacteria. These systems were also designed for use by bacteria that readily colonize the tumor interior, where these hallmark metabolic conditions are more pronounced, and so they may not offer the same tumor specificity to WCFS1 which is not known to be tumor-penetrating. In contrast, our results indicate that our sensor strain WCFS1-sppKR did not colonize tumors, but could still detect sppIP secreted by tumor cells and respond by creating significantly increased urinary nLuc.

For this, we chose to inject bacteria directly into subcutaneous tumors, as the presence of subcutaneous tumors were easily verified before bacterial administration. However, a similar detection scheme may be possible for tumors located in the lungs. The choice to detect sppIP as a model tumor secreted peptide, though synthetic, allowed us to gauge the specificity of WCFS1-sppKR *in vivo*, by comparing the response to tumors which were identical in all but sppIP secretion. There are a number of specific peptides, such as cytokines or extracellular matrix degradation products that are known to occur at significantly elevated concentrations surrounding a tumor and might be similarly sensed by bacteria with engineered receptors in the future.

Together, these results indicate the feasibility of using engineered commensal bacteria as biosensor probes for lung cancer. This application of bacterial biosensors within the critical pulmonary interface represents an under-studied and advantageous approach. Perhaps the most exciting aspect of this study is to suggest that powerful advances in synthetic biology may now be brought to bear on detecting diseases within the lung.^31^

## Methods

### Strains and Growth Conditions

*Lactiplantibacillus plantarum* strain WCFS1 was purchased from the American Type Culture Collection (BAA-793) and was routinely grown in MRS media from Research Products International (L11000-1000.0) sterilized via filtration. WCFS1 was routinely grown in stationary culture at 37°C in a CO_2_ jacketed incubator set to 5%. *Escherichia Coli* DH5alpha or BL21-DE3 strains were purchased from New England Biolabs (C2987 and C2527, respectively) and were routinely grown in LB media (Sigma) at 37°C with constant shaking. Antibiotics used for selection were purchased from Sigma Aldrich or Gold Bio.

### Electroporation

WCFS1 was made competent for electroporation following a previously published protocol.^32^ Briefly, 1 ml of a 5 ml overnight culture of WCFS1 grown in MRS media was added to 24 ml of fresh MRS containing 3% glycine and then grown in a sealed tube with shaking at 37°C until reaching an OD_600_ of 0.8 (approximately 3 hours). Cells were then washed with 5 ml of 10mM MgCl_2_ twice and SacGly solution (10% glycerol with 0.5 M sucrose) once, by centrifuging the suspension at 4,000 g for 10mins at 4°C, removing the supernatant and resuspending the cell pellet via gently vortexing. Cells were washed one final time by resuspending in 1 ml of SacGly solution and then centrifuging at 20,000 g for 1 min, then resuspended in 500 μl of fresh SacGly solution. Then, 60 μl of this cell suspension was inserted in a 1 mm gap cuvette along with the transforming DNA and electroporated in a Bio-Rad X-cell system using 1.8 kV, 200 Ω resistance, and 25 μF capacitance settings. Electroporated cells were immediately rescued with 1 ml of MRS and recovered for 4 hours at 37°C before plating.

### Recombineering

Recombination in WCFS1 was accomplished by adapting a previously published method which uses native phage recombinases.^22,23^ First, helper shuttle plasmid pLH01 (a kind gift from Sheng Yang -Addgene plasmid # 117261), which contains *L. plantarum* specific phage recombinases under the control of a peptide inducible expression system, was electroporated into WCFS1 to create a chloramphenicol resistant based editing strain WCFS1-pLH01. Next linear, double-stranded homologous recombination templates were created containing the desired insert sequence along with an erythromycin resistance cassette and approximately 1 kb of homology up and downstream of the desired insertion site. To facilitate this construction, the insertion sequence was first sub-cloned onto the shuttle plasmid pTRKH2 adjacent to the erythromycin resistance cassette (Rodolphe Barrangou & Todd Klaenhammer -Addgene plasmid # 71312). Then the insert sequence and erythromycin resistance cassette were amplified together via PCR and assembled with up and downstream homology arms via overlap assembly PCR. PCR products were purified via agarose gel extraction. WCFS1-pLH01 was prepared for electroporation following the protocol listed above while inducing recombinase expression by adding sppIP peptide (Sequence: MAGNSSNFIHKIKQIFTHR, synthesized by Genscript) to a final concentration of 0.5 μg/ml when the MRS-glycine culture reached an OD_600_ of 0.5 -0.6. Then competent cells were electroporated with approximately 500 ng of purified recombination template and plated on MRS plates containing 10 μg/ml erythromycin. Recombinant colonies were confirmed via PCR and sanger sequencing.

### Animal Handling and Care

All mice were C57BL/6J wildtype and purchased from Jackson Labs (strain 000664). All experiments were approved by the Institutional Animal Care and Use Committee at Oregon Health and Science University (TR02_IP00000674). Female mice aged between 8-16 weeks were used for all experiments

…

### Pulmonary Delivery and Bioluminescence Analysis

*L. plantarum* strains were prepared for administration by diluting an overnight culture in fresh MRS with 10 μg/ml erythromycin to an OD_600_ of 0.3. This culture was grown without agitation at 37°C until it reached an optical density OD600 of 0.8, then it was centrifuged at 4,000xg for 10 mins and the resulting cell pellet was resuspended in 0.015 of the original culture volume of filter sterilized PBS. This was then introduced to mice via oropharyngeal administration (OPA) within 30 mins. Briefly, mice were anesthetized using isoflurane gas and held vertically by their incisors such that their mouths lay open. Forty microliters of resuspended bacteria were then pipetted onto the top of the trachea and mice were allowed to breath in the suspension. For in vivo luminescence imaging, 20 μl of 5 mg/ml coelenterazine-h (NanoLight Technology) was injected via tail vein and mice were immediately imaged using a biophotonics imaging system (IVIS Spectrum, Caliper Life Sciences). Coelenterazine-h injections were performed before each imaging time point and mice were returned to normal housing cages between time points. To collect urine for nLuc activity measurements, mice were scruffed over a clean surface until they urinated (usually immediately). Afterward, urine was collected from the surface via pipette.

### Fluorescence In-Situ and Microscopy

Mouse lungs were dissected and immediately placed in a fixative solution containing 4% Paraformaldehyde in PBS for 3 hours at 4°C, then immersed in O.C.T. compound (Tissue-Tek) and frozen at -80°C in sectioning molds. Using a cryotome, 12 μm thick sections were made, transferred to microscope slides and again frozen at -20°C. To prepare for staining, slides were moved to a vacuum chamber to dry for 30 mins, immersed in fix solution for an additional 15 mins, immersed in PBS for 5 mins and then treated with 10 mg/ml Lysozyme (Sigma, L6876) in 100 mM Tris-HCl (pH 8) with 50 mM EDTA for 5 mins (pipetted directly over section). Slides were then rinsed twice with water. Slides were then incubated in a humidified chamber at 40°C overnight with a probe solution consisting of 100 μM of fluorophore conjugated probe (5’-Cy5 – CCAATCAATACCAGAGTTCG) (Integrated DNA Technologies) diluted in hybridization buffer (20mM Tris-HCl (pH 8),1 M NaCl, 0.01% SDS). This probe sequence was designed to bind to bases 67-86 of the 16S rRNA specific to strain WCFS1 (locus LP_RS02320). Slides were then rinsed with 48°C hybridization buffer and then immersed in the same for 15 mins. Slides were mounted with SlowFade Diamond Antifade Mountant with DAPI (Thermo Fisher, S36973) and imaged using a Zeiss LSM 880 inverted confocal microscope.

### Nanoluciferase Standard Production

Nanoluciferase (nLuc) was recombinantly produced and purified from *E. coli*. The nLuc coding sequence was codon optimized for *E. coli* and inserted onto plasmid pET21b+ in frame with a C-terminal His-tag. *E. coli* strain BL21-DE3 was transformed with this vector via heat shock and selection with 100 μg/ml ampicillin. nLuc expression was accomplished by growing transformed *E. coli* overnight at 37°C and then diluting 1 ml of this overnight into a 100 ml expression culture in an Erlenmeyer flask. When this expression culture reached an OD_600_ of 0.4 -0.5, IPTG (Gold Bio, I2481C) was added to a final concentration of 1 mM to induce nLuc expression. After 3 hours of expression at 37°C, bacteria were harvested by centrifuging at 10,000 g for 10 mins. The cell pellet was lysed by freezing at -20°C, then resuspending in 35 ml equilibration buffer (300 mM NaCl, 10 mM imidazole in PBS) and sonicating in an ice bath with a 1 cm diameter probe (Q sonics, Q500). Insoluble material was removed by centrifuging at 12,000 g for 20 mins, then the nLuc was purified from the supernatant using Ni-NTA agarose beads (Thermo Scientific, #88221). Purification was done in batch format, using centrifugation at 700 g for 2 mins to separate beads and vortexing to resuspend them. All steps were carried out at 4°C. Beads were added to the supernatant to bind his tagged nLuc for several hours, then washed four times with 25 mM imidazole and eluted with 250 mM imidazole. Purified nLuc was analyzed for purity via SDS PAGE gel (**Supplementary Figure S3**), buffer exchanged into PBS using 7 kDa MWCO desalt spin columns (Thermo Scientific, #89882) and then measured for absorbance at 280 nm to determine its concentration. It was then frozen in aliquots at -20°C until needed, and thawed the same day as used.

### Nanoluciferase (nLuc) Measurement

nLuc activity was read by first neutralizing 3 μl of sample in 27 μl of 1 M Tris buffer (pH 8), then adding 30 μl of pre-mixed nGlo buffer with the recommended substrate concentration (Promega, #N1110). Luminescence was read in black opaque 96 well plates (Thermo Scientific) using a TECAN Spark 20M plate reader (Serial number: 1605005980) programed to orbital shake 30s, then measure luminescence within the 430-500 nm wavelength window for 1 second.

### Xylose Induction within the Lungs

WCFS1-xylR were created using the recombineering method listed above and routinely grown in MRS with 10 μg/ml erythromycin without shaking in a 5% CO_2_ incubator. Xylose induction in vitro was gauged by growing bacteria to mid-log (OD = 0.8), centrifuging at 4,000 g for 10mins, resuspending in 1/10^th^ the volume of PBS and then back diluting 1:10 in DMEM-FBS containing various concentrations of xylose in a 96-well plate. This mixture was again incubated for 3 hours, after which secreted nLuc was measured from each mixture. For induction in vivo, WCFS1-xylR was grown to an OD of 0.8 and then spun down and resuspended in 0.0075 the initial volume in PBS. Twenty microliters of this resuspension was then mixed with 20 μl of saline with or without 20% (w/v) xylose immediately before administering all 40μl to mice via OPA. Urine from these mice were collected by scruffing over a clean surface, stored at 4°C for less than 24 h and then measured for nLuc activity.

### Tumor Detection via SppKR and SppIP

LLC cells (LL/2, CRL-1642^™^) cells were purchased from the American Type Culture Collection and routinely cultured in DMEM with 10% Fetal Bovine Serum without Ca or Mg. To create sppIP secreting LLC cells we first designed a muSEAP-FCS-sppIP coding sequence using the native muSEAP mRNA sequence from accession number AY054302 fused to a furin cleavage site (TRSRKKR) and sppIP sequence (MAGNSSNFIHKIKQIFTHR), both codon-optimized for mouse expression using IDT’s codon optimization tool. This coding sequence was inserted behind the CMV promoter within the pLenti CMV/TO Puro DEST lentiviral vector (Addgene plasmid #17293, provided by Eric Campeau & Paul Kaufman), sequence verified, and then packaged in lentiviral particles using HEK293T cells and finally transduced into LLC cells. Transduced cells were further selected using 1μg/ml Puromycin for 24 h. The resulting transduced cell line (LLC-sppIP) was kept in liquid nitrogen vapor phase until needed, and routinely cultured in DMEM with 10% FBS. For in-vitro activation of WCFS1-sppKR, WCFS1 was grown to mid-log (OD = 0.8) and then spun down at 4,000 g for 10 min before resuspending in 1/10^th^ the volume in PBS. This was then back diluted 1:10 into DMEM-FBS media used for growing LLC-sppIP cells or fresh media containing synthetic sppIP peptide (Genscript). Media from LLC-sppIP cells was collected after 24 h of growth in 96 well plates.

Mice were injected subcutaneously with 5 × 10^5^ LLC or LLC-sppIP cells. After two weeks, tumors were visible. WCFS1-sppKR were grown to mid-log (OD = 0.8), centrifuged at 4,000 g for 10 min and resuspended in 0.015 the initial culture volume in sterile filtered PBS. Thirty microliters of this bacterial suspension was then injected directly into tumors and urine was collected and measured for nLuc activity as described above.

### Statistical Analysis

All statistical analysis was done using Prism 10 for macOS. Dose response curves for WCFS1-sppKR and WCFS1-xylR were fit using non-linear regression to a variable-slope Hill equation. All data used for statistical comparisons passed the Shapiro-Wilk test for normality (P > 0.05). Students two-sided t-test was used for all statistical comparisons and an alpha of P < 0.05 was used to define significance. nLuc signals (in RLU) from mouse urine were log transformed before analysis, and charted as shown.

## Acknowledgements

This project was supported by funding (Full7340220, Full 2021-1439) from the Cancer Early Detection Advanced Research Center at Oregon Health & Science University, Knight Cancer Institute (Project Lead: Michael Brasino). Additional support was awarded from the Collins Medical Trust (ACNCR1235, PI: Michael Brasino) and the NIH (NIH/NIDCR DE028252, PI: Justin Merritt). We would like to thank Dr. Michelle Barton for guidance and insight throughout this project and Dr. Sean Speese for assistance with light microscopy. We would also like to thank Dr. Sheng Yang for providing the plasmid pLH01.

## Notes

### Competing Interest Statement

The authors have declared no competing interest.

## References

1. Escobar, L. E., Molina-Cruz, A. & Barillas-Mury, C. BCG vaccine protection from severe coronavirus disease 2019 (COVID-19). Proc. Natl. Acad. Sci. 117, 17720–17726 (2020).

2. Forbes, N. S. Engineering the perfect (bacterial) cancer therapy. Nat. Rev. Cancer 10, 785–794 (2010).

3. Charbonneau, M. R., Isabella, V. M., Li, N. & Kurtz, C. B. Developing a new class of engineered live bacterial therapeutics to treat human diseases. Nat. Commun. 11, 1738 (2020).

4. Riglar, D. T. & Silver, P. A. Engineering bacteria for diagnostic and therapeutic applications. Nat. Rev. Microbiol. (2018) doi:10.1038/nrmicro.2017.172.

5. Bourdeau, R. W. et al. Acoustic reporter genes for noninvasive imaging of microorganisms in mammalian hosts. Nature 553, 86 (2018).

6. Hurt, R. C. et al. Genomically mined acoustic reporter genes for real-time in vivo monitoring of tumors and tumor-homing bacteria. Nat. Biotechnol. 1–13 (2023) doi:10.1038/s41587-022-01581-y.

7. Danino, T. et al. Programmable probiotics for detection of cancer in urine. Sci. Transl. Med. 7, 289ra84–289ra84 (2015).

8. Riglar, D. T. et al. Engineered bacteria can function in the mammalian gut long-term as live diagnostics of inflammation. Nat. Biotechnol. 35, 653–658 (2017).

9. Mazzolini, R. et al. Engineered live bacteria suppress Pseudomonas aeruginosa infection in mouse lung and dissolve endotracheal-tube biofilms. Nat. Biotechnol. 1–10 (2023) doi:10.1038/s41587-022-01584-9.

10. Siegel, R. L., Miller, K. D., Wagle, N. S. & Jemal, A. Cancer statistics, 2023. CA. Cancer J. Clin. 73, 17–48 (2023).

11. Eberth, J. M. Geographic Availability of Low-Dose Computed Tomography for Lung Cancer Screening in the United States, 2017. Prev. Chronic. Dis. 15, (2018).

12. Zhou, N. et al. The majority of patients with resectable incidental lung cancers are ineligible for lung cancer screening. JTCVS Open 13, 379–388 (2022).

13. Yong, P. C., Sigel, K., Rehmani, S., Wisnivesky, J. & Kale, M. S. Lung Cancer Screening Uptake in the United States. Chest 157, 236–238 (2020).

14. Kleerebezem, M. et al. Complete genome sequence of Lactobacillus plantarum WCFS1. Proc. Natl. Acad. Sci. 100, 1990–1995 (2003).

15. van den Nieuwboer, M., van Hemert, S., Claassen, E. & de Vos, W. M. Lactobacillus plantarum WCFS1 and its host interaction: a dozen years after the genome. Microb. Biotechnol. 9, 452–465 (2016).

16. O’Sullivan, D. J. & Klaenhammer, T. R. High- and low-copy-number Lactococcus shuttle cloning vectors with features for clone screening. Gene 137, 227–231 (1993).

17. Karlskås, I. L. et al. Heterologous Protein Secretion in Lactobacilli with Modified pSIP Vectors. PLoS ONE 9, (2014).

18. Mathiesen, G. et al. Genome-wide analysis of signal peptide functionality in Lactobacillus plantarum WCFS1. BMC Genomics 10, 425 (2009).

19. Mathiesen, G., Sveen, A., Piard, J.-C., Axelsson, L. & Eijsink, V. G. H. Heterologous protein secretion by Lactobacillus plantarum using homologous signal peptides. J. Appl. Microbiol. 105, 215–226 (2008).

20. Spangler, J. R., Caruana, J. C., Phillips, D. A. & Walper, S. A. Broad range shuttle vector construction and promoter evaluation for the use of Lactobacillus plantarum WCFS1 as a microbial engineering platform. Synth. Biol. 4, (2019).

21. Rud, I., Jensen, P. R., Naterstad, K. & Axelsson, L. A synthetic promoter library for constitutive gene expression in Lactobacillus plantarum. Microbiology, 152, 1011–1019 (2006).

22. Huang, H., Song, X. & Yang, S. Development of a RecE/T-Assisted CRISPR–Cas9 Toolbox for Lactobacillus. Biotechnol. J. 14, 1800690 (2019).

23. Yang, P., Wang, J. & Qi, Q. Prophage recombinases-mediated genome engineering in Lactobacillus plantarum. Microb. Cell Factories 14, 154 (2015).

24. Le Noci, V. et al. Modulation of Pulmonary Microbiota by Antibiotic or Probiotic Aerosol Therapy: A Strategy to Promote Immunosurveillance against Lung Metastases. Cell Rep. 24, 3528–3538 (2018).

25. Merritt, J., Senpuku, H. & Kreth, J. Let there be bioluminescence – Development of a biophotonic imaging platform for in situ analyses of oral biofilms in animal models. Environ. Microbiol. 18, 174–190 (2016).

26. Hall, M. P. et al. Engineered Luciferase Reporter from a Deep Sea Shrimp Utilizing a Novel Imidazopyrazinone Substrate. ACS Chem. Biol. 7, 1848–1857 (2012).

27. Sasikumar, P., Gomathi, S., Anbazhagan, K. & Selvam, G. S. Secretion of Biologically Active Heterologous Oxalate Decarboxylase (OxdC) in Lactobacillus plantarum WCFS1 Using Homologous Signal Peptides. BioMed Research International https://www.hindawi.com/journals/bmri/2013/280432/ (2013) xdoi:10.1155/2013/280432.

28. Kotula, J. W. et al. Programmable bacteria detect and record an environmental signal in the mammalian gut. Proc. Natl. Acad. Sci. U. S. A. 111, 4838–4843 (2014).

29. Schellenberger, V. et al. A recombinant polypeptide extends the in vivo half-life of peptides and proteins in a tunable manner. Nat. Biotechnol. 27, 1186–1190 (2009).

30. Chien, T. et al. Enhancing the tropism of bacteria via genetically programmed biosensors. Nat. Biomed. Eng. 6, 94–104 (2022).

31. Cooper, R. M. et al. Engineered bacteria detect tumor DNA. Science 381, 682–686 (2023).

32. Leenay, R. T. et al. Genome Editing with CRISPR-Cas9 in Lactobacillus plantarum Revealed That Editing Outcomes Can Vary Across Strains and Between Methods. Biotechnol. J. 14, 1700583 (2019).

