## Supplemental Figures for "Engineered Lactiplantibacillus plantarum as a Biosensor Probe for the Lungs"

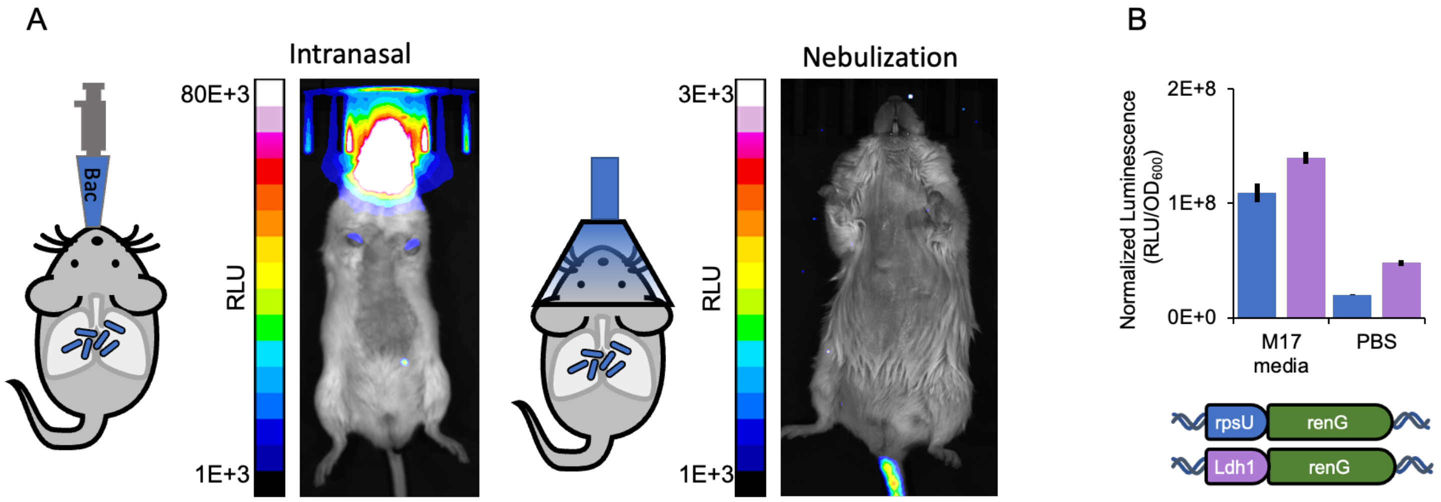


**Supplemental Figure S1: Alternative delivery methods and Comparison of rpsU and ldh1 renG luminescence.**
(**A**) IVIS images from alternative delivery methods. (**B)** WCFS1-ldh1-renG (purple) and WCFS1-rpsU-renG (blue) were grown overnight in a 37°C incubator under a 5% $CO_2$ atmosphere in MRS media, then diluted in 2-fold serial dilutions in the indicated media and incubated an additional 3 hours. Coelenterazine-h was added to each bacterial dilution to a final concentration of 4.1 μM and then luminescence and $OD_{600}$ was read in a TECAN plate reader in a 96 well plate. Bar heights indicate the average signal of n=3 dilutions with error bars showing S.E.M.

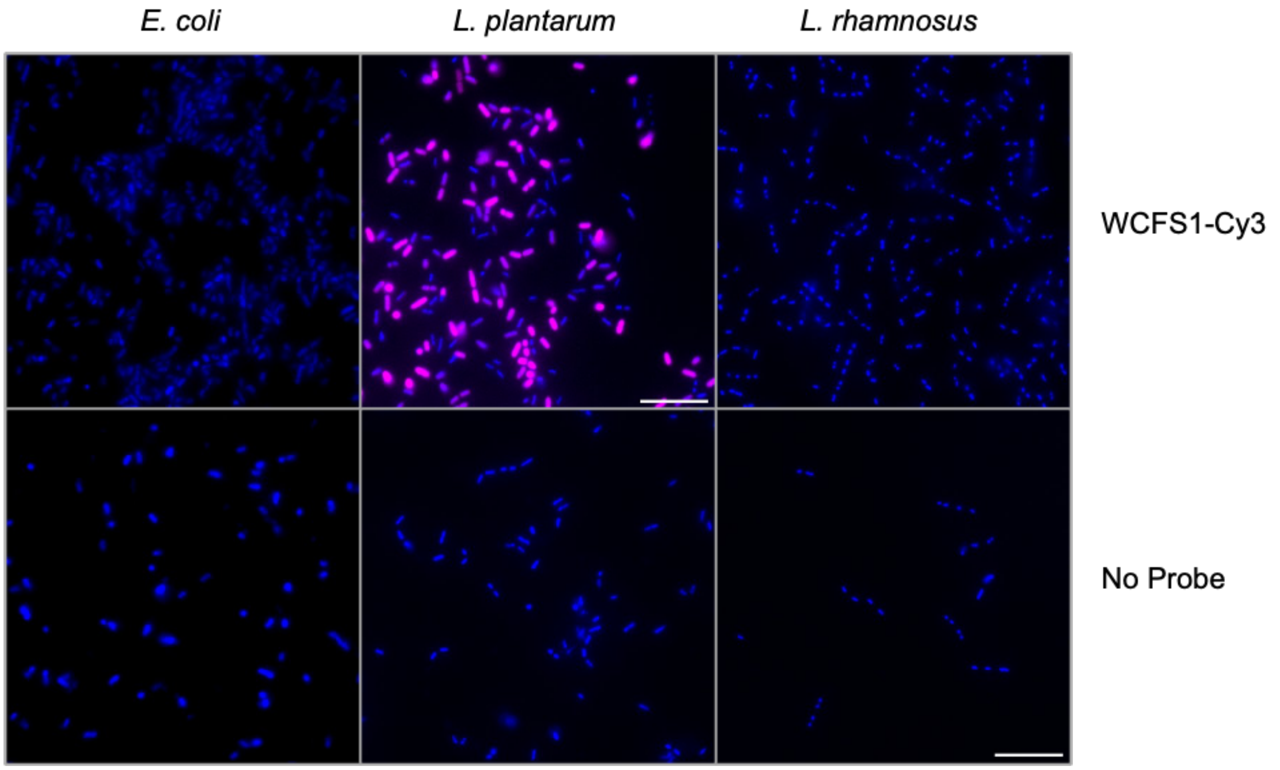


**Supplemental Figure S2: Fish Probe Specificity.** The bacterial cells shown were spun down from stationary phase and fixed on gelatin coated slides. Slides were then subjected to the same hybridization protocol as in Figure 1D. Strains used are *Escherichia coli* NIssle 1917, *L. plantarum* WCFS1 and *Lacticaseibacillus rhamnosus* GG. Look up tables for all images are identical, with DAPI signal in blue and Cy3 in magenta. All images are at thes same scale and scale bars are 10 μm.

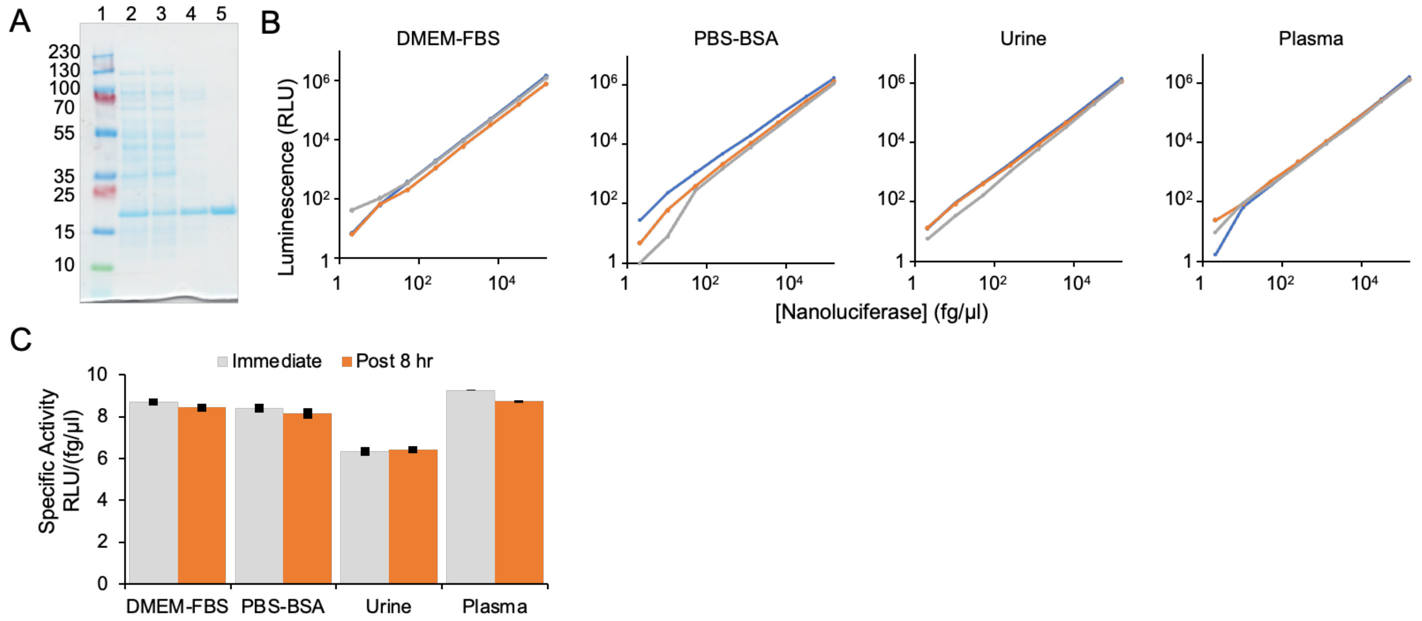


**Supplemental Figure S3: nLuc stability and activity in various media (A)** Polyacrylamide gel electrophoresis analysis of his-tagged nanoluciferase purification from *E. coli.* (1) PageRuler Plus pre-stained protein ladder (ThermoFisher, #26619), (2) soluble fraction of lysate, (3) Flow through post Ni-NTA agarose bead binding, (4) Wash, (5) Elution. **(B)** Nanoluciferase **(**nLuc) remained active and stable when diluted into various biological media. nLuc purified from *E. coli* was diluted in defined media (PBS with 10mg/ml BSA), complete cell media (DMEM with 10% v/v FBS), or urine or plasma pooled from several mice and read for activity. Immediately before each reading, samples were neutralized by 1:10 dilution in 1M Tris buffer (pH8) and then mixed 1:1 with complete nGlo buffer (Promega) per the manufacturers instructions. Readings from three independent serial dilutions are shown. The average specific activity over all dilutions was similar for all media. Luminescence was linear with nLuc concentration in all media ($R > 0.999$). **(C)** Purified nLuc was diluted in the same media as (B) from 160 pg/µl to 1.28 pg/µl and either measured for luminescence immediately or after incubating for 8 hours at 4°C. There was no significant difference between luminescence readings from DMEM, PBS or urine samples and only a slight drop when incubated in plasma, likely from protease degradation. Data represent average of n=4 dilutions, error bars are S.E.M. **(C)** Polyacrylamide gel electrophoresis analysis of his-tagged nanoluciferase purification from *E. coli.* (1) PageRuler Plus pre-stained protein ladder (ThermoFisher, #26619), (2) soluble fraction of lysate, (3) Flow through post Ni-NTA agarose bead binding, (4) Wash, (5) Elution.

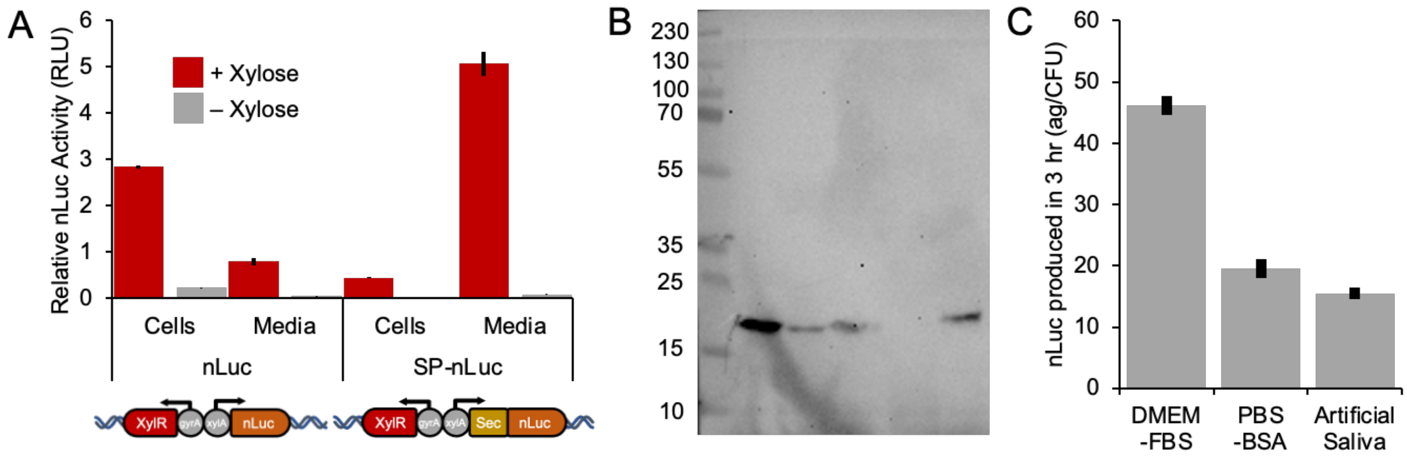


**Supplemental Figure S4: Function of nLuc secretion tag in WCFS1 (A)** WCFS1 was transformed with a pTRKH2 based plasmid carrying a codon optimized nanoluciferase alone (nLuc) or with a secretion signal peptide (SP-nLuc) taken from WCFS1 gene Lp-3050 (RefSeq WP_011102021), both under the control of a xylose inducible promoter. Relative nLuc activity associated with the cell pellet or spent growth media was measured from stationary phase overnight cultures grown with (red) or without (grey) 1% w/v xylose induction agent. Data represent average of n=2 technical replicates, error bars are S.E.M. **(B)** Composite image of membrane western-blotted with anti-nLuc primary antibody (R&D Systems, MAB#10026, Clone # 965808) and HRP conjugated secondary antibody. Composite shows chemiluminescence for blots and photo for pre-stained ladder. Pre-stained protein ladder is in the left most lane. The next 3 lanes are supernatant from rpsU-nLuc grown in (from left to right) MRS, PBS with 10% BSA or Artificial Saliva, concentrated 10x using a 10 kDa MW cut off spin filter. The next lane is blank, followed by 5ng of his-tagged nLuc purified from *E. coli*. **(C)** WCFS1-rpsU-nLuc was grown to mid-log (OD = 0.8) in rich media (MRS) and then spun down and resuspended in the indicated media in 5x serial dilutions. Agar plates were used to measure the CFUs added to each mixture. After incubating for 3 hour at 37°C in an incubator with 5% $CO_2$ atmosphere, the media was collected and measured for nLuc activity. nLuc activity was compared to purified nLuc controls to determine the final concentration. Bar height indicates the average values from 3 separate bacterial cultures, and 4 serial dilutions each, for a total of 12 data points. Error bars indicate the S.E.M. We did not observe any trend in fg/CFU with bacterial concentration in any of the media tested.

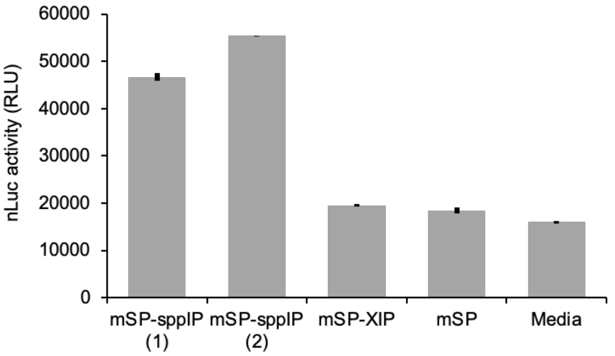


**Supplemental Figure S5**: **Specificity of the museap-sppIP to sppKR system.** Media from LLC cells were transduced with the expression construct for murine SEAP fused via a furin cleavage sequence to the sppIP peptide (mSP-sppIP), or a different bacterial peptide with sequence LPYFAGCL (mSP-XIP) or no peptide (mSP). After selecting with puromycin, media from transduced cells was incubated with mid-log WCFS1-sppKR for 3 h and the resulting nLuc activity in the media is shown (N=3, error bars are S.E.M.). Results are shown for two independent transductions of the mSP-sppIP construct as well as for fresh media.

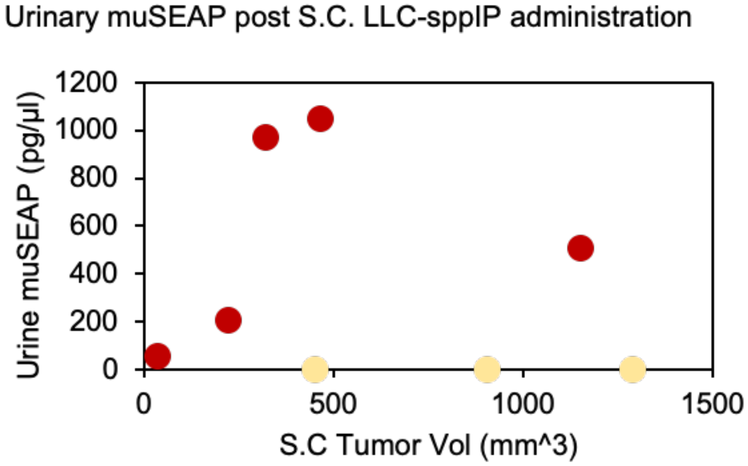


**Supplemental Figure S6**: **Tumor size vs urinary muSEAP levels.** Tumor volume was calculated by measuring tumor diameter via calipers and approximating each tumor as a sphere. All tumors appeared roughly spherical. The calculated tumor volume is plotted against muSEAP activity in urine collected 2 weeks after tumor cell injection, but before bacterial injections.
